# Hyphal differentiation and cell wall remodeling in *Trametes versicolor* underlie a layered architecture in pure mycelium materials

**DOI:** 10.64898/2026.11.19.765702

**Authors:** Annah-Ololade Sangosanya, Antoine Loquet, Agathe Ecoutin, Sofie Verschraegen, Phebe Arnauts, Joost Brancart, Niko Van den Brande, Lode Daelemans, Karen De Clerck, Mario Amalfi, Eveline Peeters

**Affiliations:** Research Group of Microbiology, Department of Bioengineering Sciences, Vrije Universiteit Brussel, Pleinlaan 2, B-1050 Brussels, Belgium; Université Bordeaux, CNRS, Bordeaux INP, CBMN, UMR 5248, IECB, F-33600 Pessac, France; Centre for Textile Science and Engineering (CTSE), Department of Materials, Textiles and Chemical Engineering (MaTCh), Ghent University, B-9052 Zwijnaarde (Ghent), Belgium; Sustainable Materials Engineering, Vrije Universiteit Brussel, Pleinlaan 2, B-1050 Brussels, Belgium; Meise Botanic Garden, Research Department, Nieuwlaan 32, B-1860 Meise, Belgium; Fédération Wallonie-Bruxelles, Service Général de l’Enseignement Supérieur et de la Recherche Scientifique, Brussels, Belgium

**Keywords:** mycelium, leather-like materials, liquid-state surface fermentation, hyphal systems, Polyporales, Basidiomycota

## Abstract

**Background:** Pure mycelium materials grown at the air–liquid interface are considered promising leather alternatives, yet they are generally treated as homogeneous sheets of fungal biomass. Because basidiomycetes differentiate their hyphae into generative, skeletal and ligative cell types, the properties of such materials may instead arise from an internal tissue organization. In this study, cell types produced by the trimitic white-rot fungus *Trametes versicolor* during liquid-state surface fermentation were characterized and the impact of their spatial organization, mechanics and cell wall composition on the resulting material was analyzed.

**Results:** Optic and electron microscopy revealed that the mats are not homogeneous but consist of two layers: a dense bottom layer of thin-walled generative hyphae in contact with the liquid medium, and a looser, air-exposed top layer made of thick-walled skeletal hyphae. Both layers displayed comparable hyphal diameters (2.43 ± 0.42 and 2.33 ± 0.50 µm), although the skeletal hyphae in the mat measured approximately half the diameter of their fruiting-body counterparts (5.10 ± 1.05 µm). Stress–strain curves of untreated mats displayed a sequential failure consisting of a brittle fracture of the generative bottom layer followed by a ductile failure of the skeletal layer. The two faces also differed in wettability, the top being hydrophobic and highly adhesive and the bottom hydrophilic. Solid-state nuclear magnetic resonance showed extensive rigid cell wall remodeling between the two cell types: α-1,3-glucan accounted for about one third of the rigid polysaccharides in the generative layer but was undetectable in the skeletal layer, which contained approximately 74% β-1,3-glucan and 26% chitin and was overall more rigid and more lipid-rich.

**Conclusions:** This work shows that *T. versicolor* differentiates into a very specific type of skeletal hyphae in liquid state surface fermentation, which involves remodeling of the molecular architecture of the cell wall rather than a simple thickening of it, and that the mechanical and surface behavior of the material can be traced to that remodeling and to the cellular organization.

**Graphical abstract:** 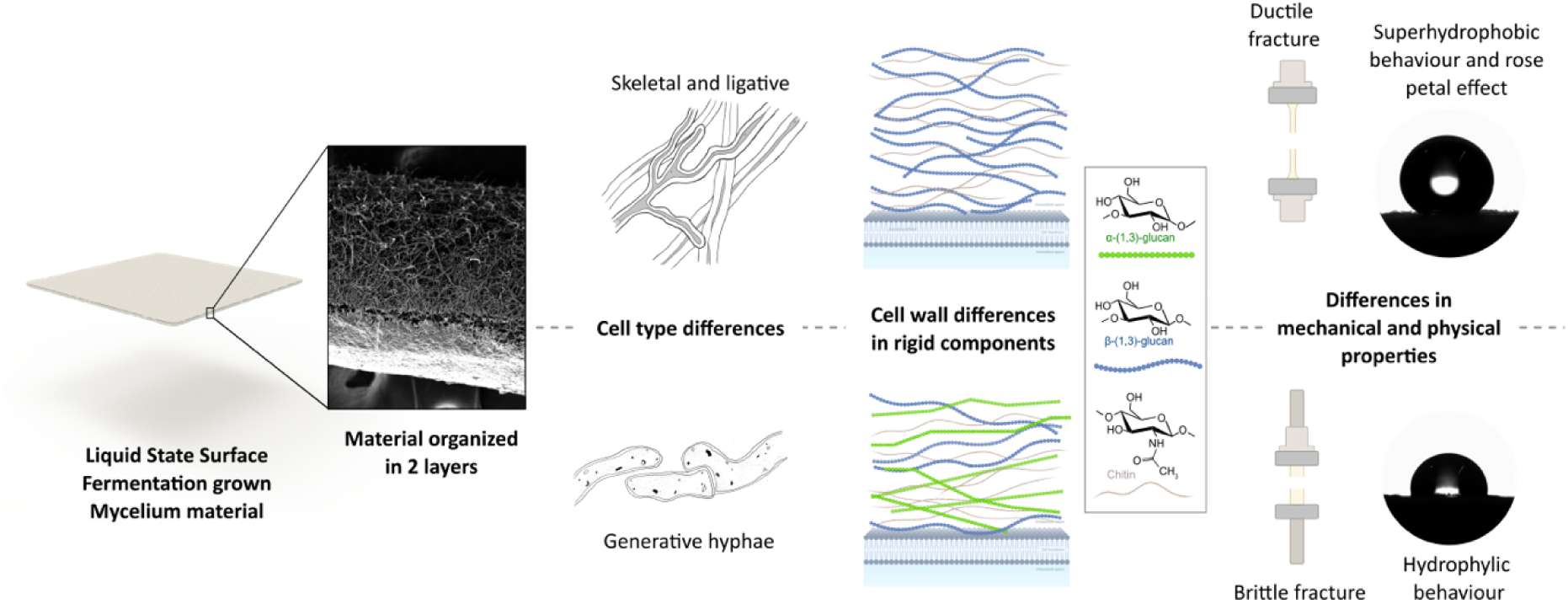

## Background

*Trametes versicolor* is a white-rot polypore fungus belonging to Basidiomycota, which is widely distributed in the North temperate regions of Europe, north America and Asia. It is recognizable from its fruiting body featuring characteristic concentric multicolored bands similar to the tail feathers of turkeys, giving it its nickname “Turkey’s tail fungus” (Lodi et al., 2025; Ryvarden et al., 2022; Ryvarden and Melo, 2014).

Despite a fruiting body of relatively small size, its biomass is valorized in various applications. Noticeably used in medicine for its β-glucans, it produces polysaccharide-K (PSK or Krestin) and polysaccharopeptide (PSP), biologically active polysaccharides associated with immunomodulation (Habtemariam, 2020; Lodi et al., 2025). Beyond medicine, *T. versicolor* living and dead biomass can be used for the biosorption of pollutants from contaminated water and industrial effluents. The fungal cell wall of *T. versicolor* contains numerous functional groups within its constituents chitin, glucans, proteins, and polysaccharides, which are capable of interacting with pollutants and absorbing them to the cell surface (Elgarahy et al., 2021). The fungal species has particularly been used in the adsorption of synthetic textile dyes (reactive azo dyes) and heavy metals (including copper, lead, zinc, cadmium, chromium…) by binding them to negatively charged functional groups from the fungal cell wall (Bayramoglu, 2003; Boujelben et al., 2024; Cengiz et al., 2014; Yalçınkaya et al., 2002). While both living and dead fungal biomass can be used as biosorbents, dead biomass is often preferred industrially because it does not require nutrients, tolerates toxic conditions better, can be regenerated and avoids complications related to fungal metabolism.

In recent years, mycelium from filamentous fungi such as *T. versicolor* has been investigated for production of sustainable biomaterials. Tuning the fermentation method, it is used to produce versatile mycelium materials. Grown on solid substrates, it can form composite materials relevant in architecture (bricks, insulation), but also for fashion (flexible textile-mycelium composites) (Gandia et al., 2021; Romero-Cedillo et al., 2025; Silm et al., 2026). An alternative strategy for producing mycelium materials is to cultivate the fungus at the air-liquid interface in a static liquid-state fermentation, where it forms a continuous pure mycelium sheet that can be readily harvested without disrupting either surface of the material (Cartabia et al., 2021; Romero-Cedillo et al., 2025), making the liquid-state surface fermentation (LSSF) set-up a highly relevant technique in an industrial context. More broadly, pure mycelium materials (PMMs) are increasingly being recognized as a new class of biofabricated materials, which properties can be tailored for applications extending well beyond leather replacement (Jones et al., 2021).

At a microscopic level, PMMs consist exclusively of fungal hyphae. Hyphae can differentiate into distinct cell types that collectively define the hyphal system of the fungus. In 1933, Corner introduced the hyphal systems classification of basidiomycetes in three categories according to the cell types produced in their basidiocarp (Corner, 1932a, 1932b; Ryvarden and Melo, 2014). Depending on the species and on the tissue, the fungus can produce generative, skeletal or ligative (connective) hyphae. Generative hyphae constitute the precursor of all other cell types and can be produced by all basidiomycetes. These cells are characterized by a thinner cell wall and the presence of septa and, in many species, clamp connections. Skeletal hyphae are longer and thick cell walled to solid, typically featuring no septa nor clamp connections (Ryvarden and Melo, 2014). It has been demonstrated that skeletal cells provide most of the mechanical strength of the basidiocarp, with their contribution estimated at 71% (Porter and Naleway, 2022). Ligative hyphae, also called binding hyphae, are typically branched, thick-walled to solid and nonseptate; they bind other hyphae together.

*T. versicolor* is classified as a trimitic strain widely used in mycelium materials (Ryvarden and Melo, 2014; Vandelook et al., 2021). It is described to produce thin-walled, hyaline generative hyphae of about 2.5-3 μm in diameter, thick-walled hyaline skeletal hyphae of about 4-8 μm in diameter as well as thick-walled, nonseptate, much-branched binding hyphae of about 2-4 μm in diameter (Ryvarden and Melo, 2014). Despite the widespread use of *T. versicolor* in medicine, bioremediation and for production of mycelium-based materials, little is known about the specific roles that such different hyphal cell types play in these applications. While a few studies have examined the influence of hyphal morphology and cell wall architecture on the properties of mycelium materials in other fungal species and in solid composite setups (Aiduang et al., 2024; Aquino et al., 2022; Gallagher et al., 2025), the contribution of individual hyphal types to the organization and properties of *T. versicolor* PMMs remains unexplored.

In this study, we characterized the cell types produced during the growth of *T. versicolor* PMMs by LSSF and investigated how their spatial organization and cell wall composition influence the properties of the resulting material.

## Methods

### Strains and cultivation conditions

*T. versicolor* VUB-SV1 (Vandelook et al., 2024) was used for all cultures. The strain was maintained on Complete Yeast Medium Agar (CYM-agar) plates containing 20 g/L glucose, 3 g/L peptone, 2 g/L yeast extract, 0.5 g/L MgSO_4_, 2.5 mL/L phosphate buffer (7.9 g/L NaCl, 0.2 g/L KCl, 3.58 g/L Na_2_HPO_4_.12H_2_O and 0.246 g/L H_2_KPO_4_) and 10 g/L micro agar. The pH was adjusted to 7.2. For liquid cultures, CYM was used without agar. Glucose, phosphate buffer and the remaining CYM medium components were autoclaved in separate bottles for 15 minutes at 121°C. All medium components were purchased from Sigma-Aldrich.

### Liquid-state surface fermentation for production of mycelium mats

Mycelium mats were grown using LSSF, a static fermentation technique. 250 mL Erlenmeyer’s with 8 cm ø base and 5.5 cm ø opening (for ease of biomass harvest) were filled with 100 mL CYM and inoculated with a 5 mm ø agar plug of *T. versicolor* grown on CYM-agar. Inoculated Erlenmeyer flasks were incubated statically in the dark for 7 days at 25°C. After 7 days, the culture was blended using a Waring 7012S 7-Speed Blender (Waring Commercial, Stamford, Connecticut, USA) and a 360 mL Stainless Steel Semi-Micro Blending Container with Press Fit Lid (Eberbach corporation, Van Buren Charter Township, MI, USA) at power 5 for 10 seconds. 5mL of this blended inoculum were used to inoculate the cultures composed of 100 mL CYM in a microbox container type O118/50+OD118 (SacO2, Deinze, Belgium). Cultures were cultivated statically in the dark for 15 days at 25°C.

Mycelium mats were harvested from their box and prior to drying, a piece of the mat was cut out (about 1 cm²). In this sample, bottom cells were manually separated from top cells using a scalpel. This operation was performed in biological triplicates. Top cells were pooled together in a same tube, and similarly bottom cells pooled together, samples were frozen for downstream nuclear magnetic resonance (NMR) analysis. A similar dissection was performed for dynamic vapor sorption (DVS) analysis samples, performed in technical triplicates and biological triplicates. These samples were placed in a 1.5 mL Eppendorf tube each with the lid open. The remainder of the mycelium mat was placed on a silicone surface to avoid stickiness. Both the mycelium mat and the DVS samples were dried in a Memmert UN110 oven at 40°C, 100% fan and 100% flap for at least 24 hours.

### Production of mycelial pellet samples

Mycelial pellets of *T. versicolor* were produced for microscopy and NMR analysis. A 250 mL Erlenmeyer with 8 cm ø base and 5.5 cm ø wide opening (for ease of biomass harvest) was filled with 100 mL CYM and inoculated with a 5-mm ø agar plug of *T. versicolor* grown on CYM-agar. After 7 days, the culture was blended using a Waring 7012S 7-Speed Blender (Waring Commercial, Stamford, Connecticut, USA) and a 360 ml Stainless Steel Semi-Micro Blending Container with Press Fit Lid (Eberbach corporation, Van Buren Charter Township, MI, USA) at power 5 for 10 seconds and was determined to be concentrated at ∼7 mg dry biomass/mL. One mL of this mycelium suspension was used to inoculate a 250 mL Erlenmeyer flask filled with 100 mL CYM and incubated while shaking at 180 rpm, 25°C for 7 days. Mycelium pellets formed were then immediately visualized under the microscope or frozen for NMR analysis.

### Preparation of fruiting body samples

A herbarium sample of *T. versicolor* was prepared from a fruiting body sampled in Saint-Vincent Park (50.88131, 4.40315) in the municipality of Evere, Belgium. The sample was dried for 24 hours in open air and subsequently frozen for 14 days to kill any parasite in the sporocarp. The sample was then dried again in open air. The sample was confirmed to be a *T. versicolor* strain by performing a Polymeric Chain Reaction (PCR) of the internal transcribed spacer region of ribosomal DNA using primers ITS1 and ITS4 (White et al., 1990) and partial Transcription Elongation Factor gene (TEF-1α) using primers EF1-983 and EF1-2218 (Brandon Matheny et al., 2007; Rehner and Buckley, 2005). The fruiting body sample was dissected using a scalpel and tweezers under a binocular microscope. Separation was made dry allowing to get a crust sample, a sample of the context and a sample of the spore tubes. Samples were placed in 1.5 mL Eppendorf tubes, rehydrated with 500 µL water, frozen and subjected to NMR analysis.

### Optical microscopy

Microscopy slides were obtained by embedding the sample into 4% agar and slicing between 30 µm and 50 µm thick slices using a VT1000S vibratome (Leica Biosystems, Nussloch, Germany) in a Phosphate Buffer Saline bath (Sigma-Aldrich, P4417 50TAB) on frequency 4, speed 5. Alternatively, slides were made by sampling cells using tweezers, addition of 5% KOH, brief flaming of the slide, and spreading the cells by addition of coverslip. Observations were carried out on an Eclipse Ti2 inverted epi-fluorescence microscope (Nikon, Tokyo, Japan) on bright field setting. Image capture and measurements were performed using the NIS-Elements BR software. Cell diameter and cell wall thickness measurements were performed by using at least 3 images and performing measurement on at least 10 random spots of the cells of interest (**Supplementary Table S1**). Cell diameters were analyzed using a linear mixed-effects model, treating experimental group as a fixed effect and image as a random effect. Significant differences between groups were identified by likelihood-ratio testing followed by Holm-adjusted pairwise comparisons.

### Scanning electron microscopy

SEM was performed using a JEOL JSM-6610LV SEM microscope equipped with Oxford X-Max detector and Aztec software (Oxford Instruments, Germany). Samples were fixed to the holder using conductive carbon tape and gold-coated to 3 nm using a Cressington 208HRD high resolution sputter Coater (Cressington Scientific Instruments, Watford,UK).

### Tensile testing

Stress strain curves were obtained using a TA Instruments RSA-G2 Dynamic Mechanical Analyzer (New Castle, DE, USA) with the Trios software. Tensile strength was measured by performing axial deformation with a Hencky strain rate of 5%/min. Sampling rate was set to 5 points/second. For each of the three biological replicates, measurements were performed in at least three technical replicates. Samples were cut into dog-bone shapes with a width of 8 mm at the narrowest point and a gauge length of 20 mm using a custom-made leather cutting die (**Supplementary Figure S1**) comprising a knife-type contour and a central foam-type pressure pad (Tools C Dies Engineering, Brecht, Belgium). Samples were placed on the clamps using a torque screwdriver at a torque of 7 cN.m. Only biological replicates with a thickness of 1 mm or higher were used for tensile strength testing.

The ultimate tensile stress (UTS) was determined from the stress-strain plots as the peak stress reached before decline of the curve. The elongation at break (EAB) was determined as the first post-peak strain at which stress fell below 5% of the UTS. Mean curves were obtained by interpolating each specimen onto a common strain axis and averaging, with each specimen contributing only up to its own break point. Technical replicates were first averaged within each biological replicate.

### Water contact angle analysis

Hydrophobicity of the samples was measured by capturing images of the water contact angle (WCA) using a dataphysics contact angle system OCA 25 equipped with a TBU100 base rotation unit. The contact angle of a 10-µL water droplet was measured over the course of 5 minutes using the SCA20 software for OCA and PCA. The baseline was set manually. Measurements were performed in five technical replicates for three biological replicates on both top and bottom sides of the mycelium mats.

Measured contact angles values were time-binned per second and the average WCA was calculated for both sides for 30 seconds and 5 minutes. Statistical differences were calculated using a two-tailed paired Student’s t-test on biological replicate means, with technical replicates averaged within each biological replicate. Droplets that were fully absorbed before the end of the acquisition, were assigned a WCA of 0°, as complete absorption represents total wetting.

Hysteresis was determined by pouring a 30-µL water droplet on the material and tilting the machine 90°C at a velocity of 0.49°/sec. Measurements were performed in three technical replicates for each of the three biological replicates. The advancing contact angle (θA, downhill/front edge angle) and the receding contact angle (θR, uphill/back edge angle) were automatically determined by the software for each tilting angle. From these angles, the hysteresis was calculated as Hysteresis = θA − θR. The roll-off angle was determined either as the machine tilting angle at which the droplet was lost from the frame or as the machine tilting angle at which the advancing angle rose above static, peaked and fell ≥15° (depinning).

For the plotting of the wetting state map, the apparent static contact angle was defined as the average of the mean contact angle of the five valid frames. The hysteresis value used was the maximum hysteresis value during the whole run (hysteresis reached before rolling off or when reaching 90°).

### Dynamic vapor sorption analysis

DVS analysis was performed to investigate the sorption/desorption behavior of the generative layer *versus* the skeletal-ligative layers of the mat. Dried samples were pre-weighted to ensure the dry weight exceeded 1 mg. Water vapor sorption and desorption characteristics were determined using a TA Instruments Q5000 SA Vapor Sorption Analyzer (New Castle, DE, USA). The dry sample mass was determined following two successive drying steps at 40°C and subsequently at 25°C under 0% relative humidity (RH). The samples were then subjected to a sorption-desorption cycle at 25 °C. RH was increased from 0% to 90% RH in 10% increments and subsequently decreased from 90% to 0% RH using the same step size. Equilibrium at each RH step was assumed when the mass change was less than 0.1% over 12 minutes, with a maximum dwell time of 300 minutes per step.

Sorption isotherms were constructed by taking the mean water uptake (% dry mass) over the final 10 minutes of each 10% RH step as that step’s equilibrium value, averaging the three technical replicates within each biological replicate, and plotting the mean across the three biological replicates.

### Solid-state nuclear magnetic resonance spectroscopy

Samples were analyzed using magic-angle spinning solid-state NMR. Samples were packed in 4 mm rotors and analyzed on a 600 MHz NMR magnet (Bruker Biospin) with a triple resonance 4 mm probe. Magic-angle spinning frequency was to 11 kHz. High power decoupling (90 kHz) was used during acquisition. Cross-polarization (CP) experiments were performed using a contact time of 1 ms. 1D CP and INEPT (Insensitive Nuclei Enhanced by Polarization Transfer) spectra were recorded with 22 k scans.

### Use of artificial intelligence tools

Generative artificial intelligence was used for debugging the Python scripts of the statistical analysis (hyphal cell diameters, tensile test curves, WCA measurements and DVS plots). All scripts were reviewed, tested and validated by the authors. GenAI was also used for language editing, limited to grammar, spelling and syntax of author-written text. The tools were not used to generate data, figures, interpretations or conclusions. The authors take full responsibility for the content, accuracy and integrity of the manuscript.

## Results

### Spatial organization of distinct hyphal types within the *T. versicolor* mycelium mat

In order to investigate the cell types produced in the mycelium mat and their organization within it, microscopy images were made using optic microscopy and SEM. The mycelium mat organization was investigated by observing a longitudinal cut, but by assessing its different areas and the observed cells were compared to the cells observable in the fruiting body (**Figure 1**).

**Figure 1.**
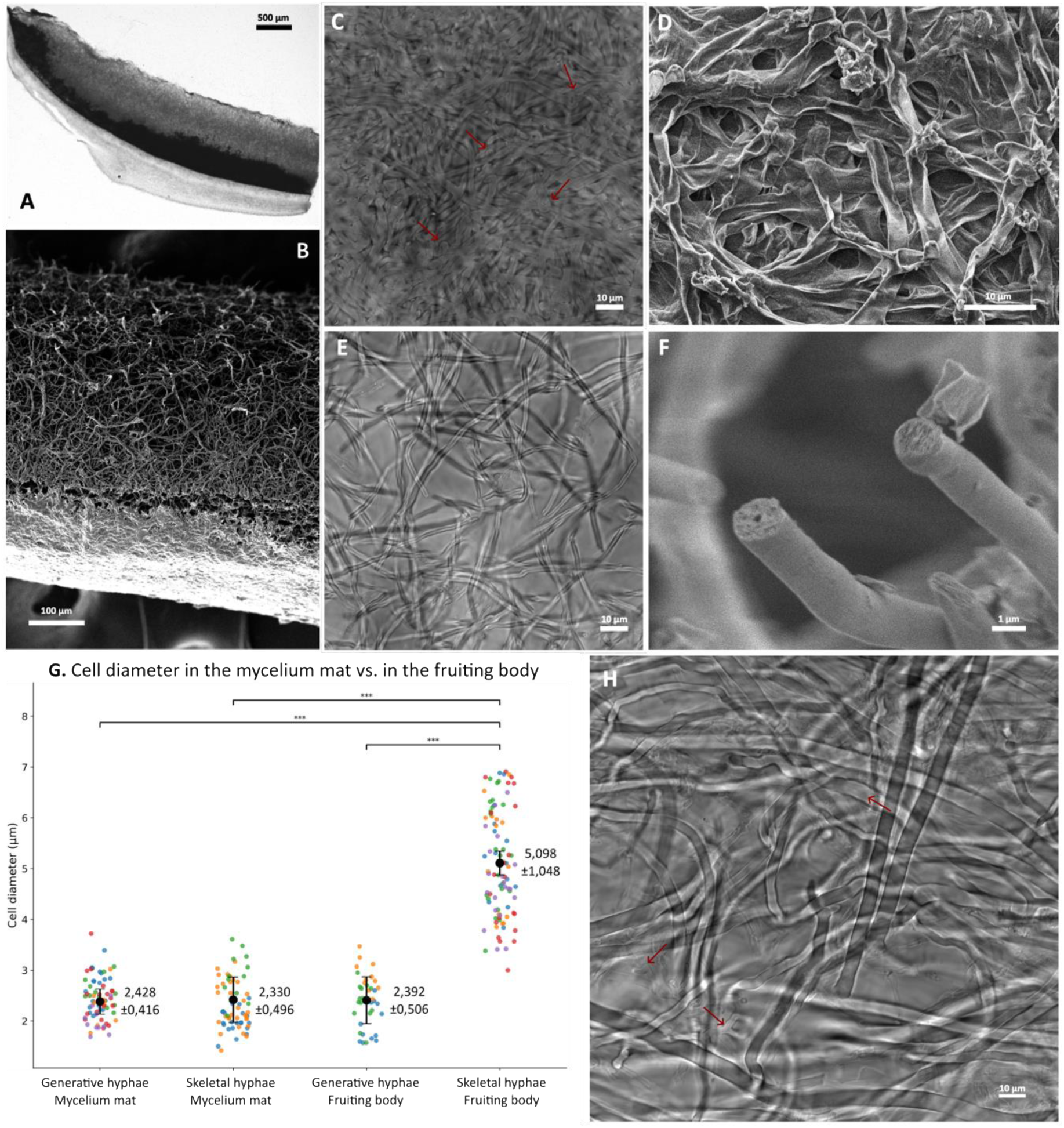
Hyphal organization and morphology in LSSF-derived mycelium mats and fruiting bodies of *T. versicolor*. **A**) Light micrograph of a cross-section of an LSSF-derived mycelium mat, with visible pigmentation line. The bottom side represents the layer in contact with the liquid medium, while the top represents the layer in contact with the air. **B**) SEM micrograph of a cross-section of an LSSF-derived mycelium mat, showing a denser layer of generative cells layer on the bottom side. **C**) Light micrograph of densely packed generative cells from the bottom layer. Visible clamp connections are indicated with red arrows. **D**) SEM micrograph of densely packed generative cells from the bottom layer, which were collapsed under the void because of their thin cell wall. **E**) Light micrograph of semi-solid skeletal cells from the top layer, showing thick cell walls. **F**) SEM micrograph of semi-solid skeletal cells from the top layer, showing thick cell walls. **G**) Dot plot of the diameters of generative and skeletal hyphal structures in the mycelium mat and in the fruiting body. Different colors represent different images used for the measurements. Brackets with *** represent a significance difference with a p-value<0.001 for the mixed linear models test. **H**) Light micrograph of hyphae from a fruiting body, with arrows indicating thicker, larger and solid skeletal hyphal cells, a thin-walled generative hyphal structure with a clamp connection and thinner, solid and branched, ligative hyphal cells, respectively.

Analysis of the mats by optic and electron microscopy revealed that the mats grown from *T. versicolor* in LSSF were not homogeneous in cellular composition. In the longitudinal cut of the sample under optic microscopy (**Figure 1A**) and SEM (**Figure 1B**), two cell layers of different densities can be distinguished. Optic microscopy also revealed the production of pigments at the interface of those two layers (**Figure 1A**). Mats were composed of a first layer of generative cells at the bottom side, which is in contact with the liquid phase during fermentation. These cells are distinguishable by their thin cell wall and clamp connections (**Figure 1C**), very densely packed and compacted together (**Figure 1D**). The average diameter of those cells was 2.428 µm ± 0.416 µm.

The second layer, in contact with the air during growth, was composed mainly of skeletal cells (**Figure 1E**), but also featured ligative and generative cells. Skeletal cells are distinguishable by their lack of clamp connections and septa and by their thick cell wall (**Figure 1F**), rendering the cells semi-solid with almost no lumen, resembling textile fibers. These cells did not differ in diameter from the generative cells with an average diameter of 2.330 µm ± 0.496 µm (**Figure 1G**).

Morphology of the different cells was also observed in a fruiting body sample as a comparison point (**Figure 1H**). Interestingly, while generative and skeletal cells from the mat had comparable diameters to each other, their diameter was significantly lower than their counterparts in the fruiting body (p-value <0.001) (**Figure 1G**). Statistical test results are presented in **Supplementary Table S2**.

Cells from a shaken liquid culture of *T. versicolor* were also visualized under the microscope. Interestingly, the cells produced in this culture setup only contained generative hyphae (**Supplementary Figure S2**).

### Mechanical properties of the mycelium mat layers

In order to understand the mechanical behavior of the material, nominal stress-strain curves were measured (**Figure 2**). Average UTS, EAB and Young’s modulus (E) values were determined from the obtained plots. UTS was determined as the highest tensile stress value reached during the test and the EAB was determined as the first post-peak point below 5% of the UTS. The Young’s modulus was determined as the slope of a least-squares linear fit to the stress–strain curve between 0.1% and 1.0% strain. On average, the samples reached a UTS of 0.40 ± 0.08 MPa, an EAB of 3.90 ± 0.13% and a Young’s modulus of 27.5 ± 4.3 MPa (**Figure 2A**, **Supplementary Table S3**).

**Figure 2.**
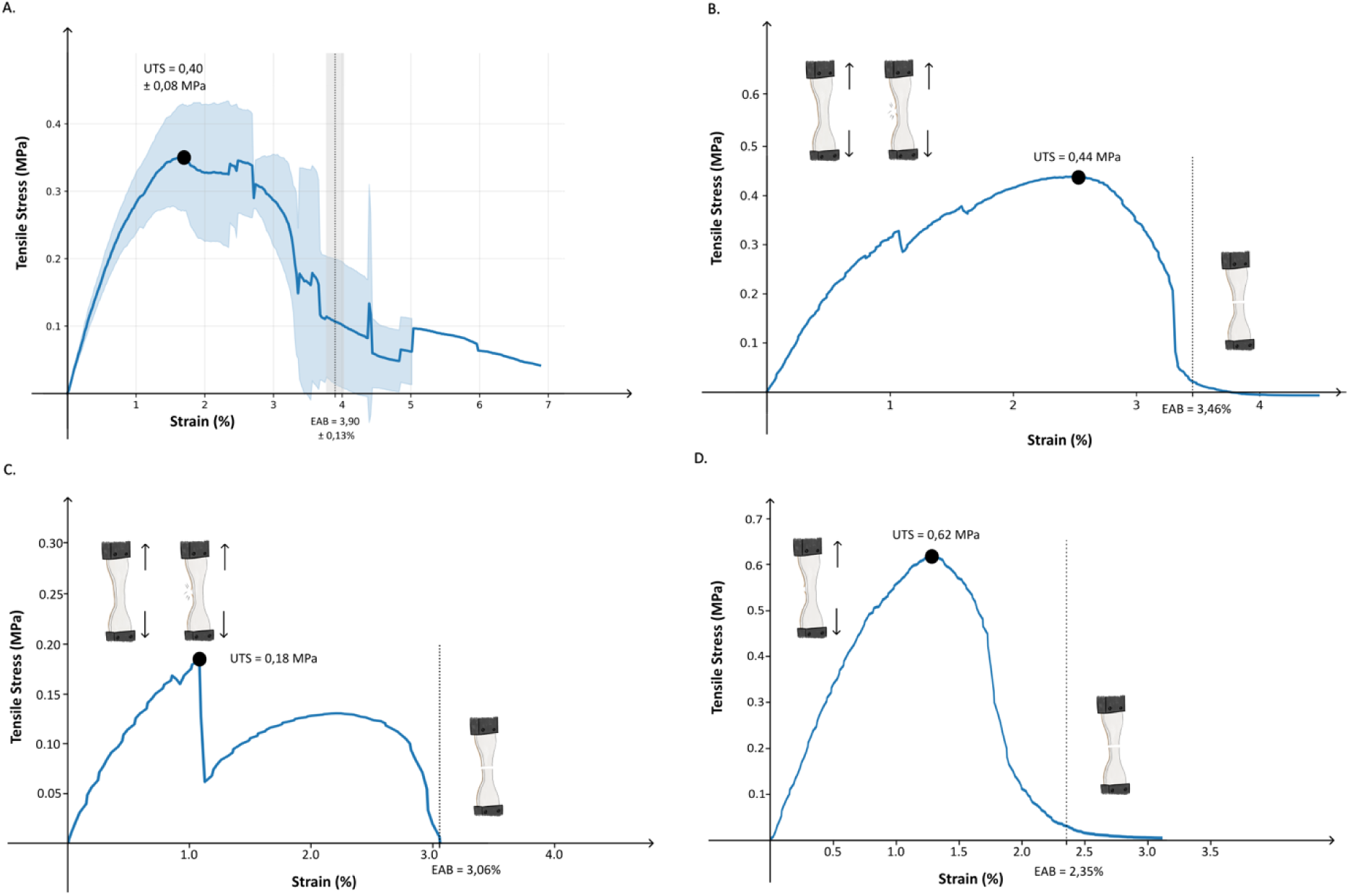
Stress-strain curves of mycelium mat samples. **A**) Average stress-strain curve. **B**)-**D**) Stress-strain curves from individual technical replicates, illustrating different fracture behaviors. **B)** The generative layer was not the load-bearing layer and cracked in two occurrences before the skeletal layer failed. **C)** The generative layer is the load-bearing layer and failed first, followed by a slower failure of the skeletal layer. **D)** The generative layer had already failed before the tensile strength test and only the skeletal layer is quantified.

Interestingly, when performing the tensile strength test, audible cracking could be heard in most samples that coincided with sudden drops in tensile stress (**Figure 2B** and **2C**). After careful observation, it appears this cracking was due to sudden breakage in the bottom layer of the mat, demonstrating that the two layers fail with different patterns (**Figure 2B-D**, **Supplementary Figure S3**, **Supplementary Video S1**).

### Surface wettability and moisture sorption of the mycelium mat layers

The WCA method was used to determine the wettability of the generative surface and of the skeletal surface (**Figure 3A**). This technique is based on the measuring of the angle that is formed between a surface and a water droplet (Rea et al., 2014). A smaller angle suggests that the water spread more easily over the surface, while a larger angle indicates lower wettability. Surfaces with a WCA below 90° are usually described as hydrophilic and those with a WCA above 90°, as hydrophobic (Rea et al., 2014).

**Figure 3.**
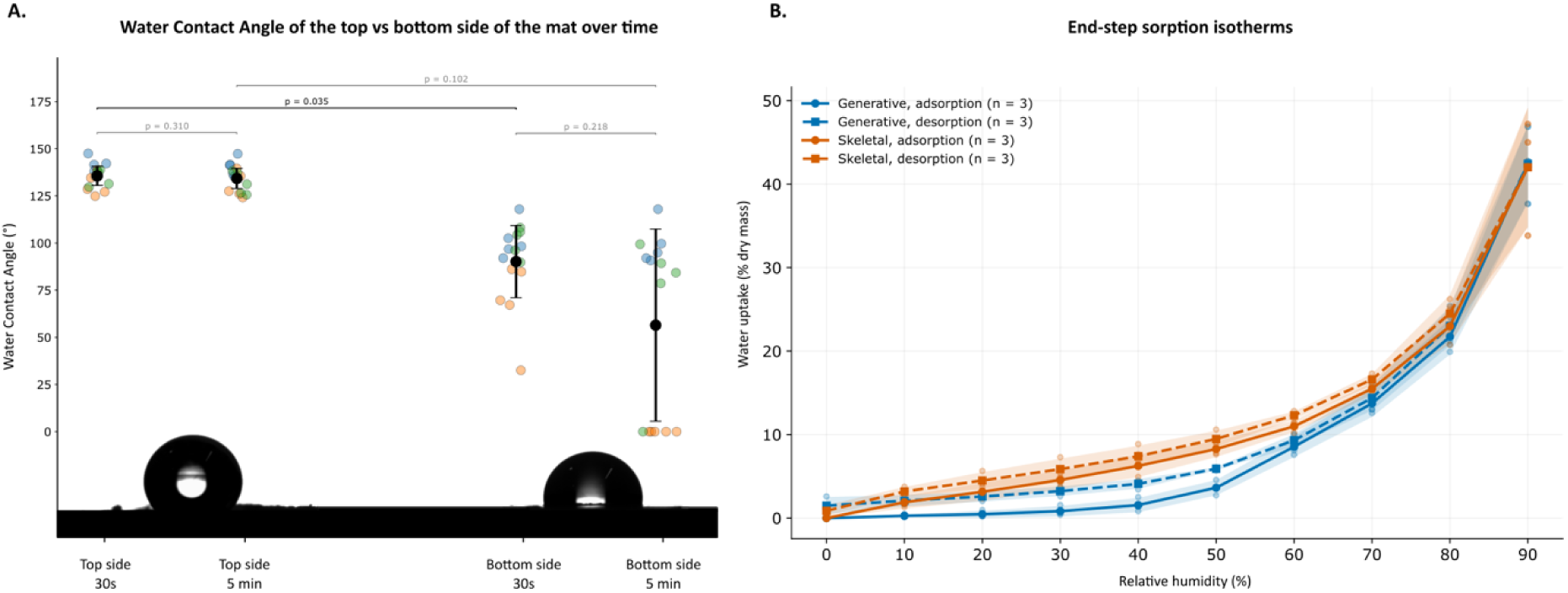
Surface wettability and moisture sorption of the two layers of the mycelium mat. **A)** WCA values of the top and bottom layers over the course of 5 minutes. Each biological replicate is represented in a different color. Significant differences are represented with a black bracket, non-significant differences with a grey bracket, with the associated p-values on top (paired t-test). **B)** DVS isotherms. Shaded bands indicate ± 1 SD across biological replicates, while individual biological means are shown as dots.

On average, the top layer featured a WCA of 135.7 ± 5.1° after 30 seconds, which did not significantly decrease after 5 minutes (134.3 ± 5.4°), indicating a constant hydrophobic/superhydrophobic behavior. The generative layer was significantly more absorbent with a WCA of 90.2 ± 19.1° after 30 seconds. After 5 minutes, 6 replicates had fully absorbed the droplet resulting in an average WCA of 56.5 ± 51.0° (**Figure 3A, Supplementary Tables S4 and S5**). Tilted WCA measurements allowed to estimate the contact angle hysteresis and the roll-off angle, as well as to establish a wetting-state map (**Supplementary Figure S4**). The results suggest that the samples exhibited a high water-droplet adhesion, with the top surface showing behavior consistent with a Cassie-impregnating or rose-petal state, characterized by high hydrophobicity and adhesion.

Subsequently, DVS analysis was performed on the generative and skeletal layers, to determine whether the observed differences surface wettability are also reflected in the moisture sorption behavior of the material (**Figure 3B**). In this technique, a sample is placed in a pan and exposed to progressively higher levels of humidity, allowing to understand the intrinsic wettability nature of each cell types.

DVS curves indicated that, at relative humidity (RH) levels below 80%, the skeletal layer sorbed more water vapor than the generative layer (**Figure 3B**). This result suggests that, despite its low surface wettability, it skeletal layer retains more humidity at lower RH levels than the generative layer. The generative isotherm remained flat until ∼30% RH, which could be explained by a limited number of accessible sorption sites, while the skeletal isotherm increased from the start. This could be explained by a looser cellular matrix, allowing for a larger surface area accessible to water vapor. This difference could also be partially explained by differences in cell wall composition. Both layers showed a steep increase from 70% RH on, which could be explained by capillary condensation. Additionally, the desorption curves were higher than the absorption curves for both layers, suggesting the mycelium mat retained water upon drying.

Interestingly, RH over most land areas typically ranges between 60% and 80% (according to Global Data Lab), which is a range at which the water uptake strongly varied for both layers, hinting that the mycelium material would be highly sensitive to climate conditions.

### The rigid cell wall composition differs between the generative- and skeletal-enriched layers

To characterize the molecular composition and dynamics of the two layers of the mycelium mat, solid-state ^13^C magic-angle spinning solid-state NMR spectra were recorded using two complementary polarization schemes. Cross-polarization (CP) preferentially detects rigid molecular components through dipolar couplings, whereas INEPT selectively detects highly mobile species through scalar couplings.

The rigid composition of the cell wall was studied first (**Figure 4)**. A comparison of the CP spectra of the bottom (generative) and top (skeletal/ligative-enriched) fractions revealed markedly different molecular fingerprints (**Figure 4A**). The most striking difference was the presence of α-1,3-glucan resonances in the bottom fraction and their complete absence from the top fraction. Consequently, the rigid polysaccharide network of the top fraction consisted exclusively of β-1,3-glucan and chitin, whereas the bottom fraction additionally contained a substantial proportion of α-1,3-glucan. Peak integration allowed to estimate the relative abundance of the major rigid polysaccharides (**Figure 4B**). The top fraction was composed of approximately 74% β-1,3-glucan and 26% chitin, whereas the bottom fraction contained approximately 26% β-1,3-glucan, 37% chitin and 37% α-1,3-glucan (**Figure 4C**).

**Figure 4.**
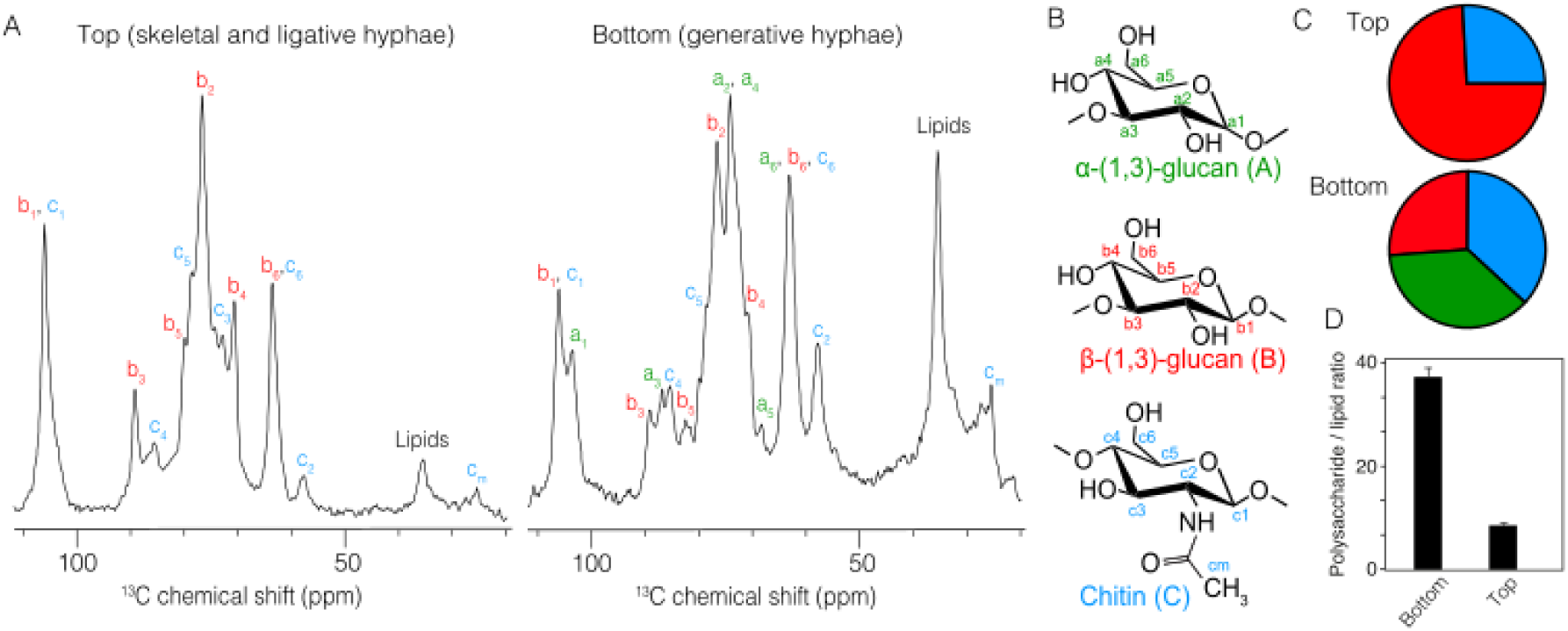
Structural analysis of the rigid component of the cell wall. **A)** Cross-polarization spectra recorded at 11 kHz MAS rate, 600MHz ^1^H frequency. α-1,3-, β-1,3-glucan and chitin are respectively assigned a, b and c with their associated carbon position and **B)** chemical structure of the three polysaccharides. **C)** Molar composition of the rigid polysaccharides, with a precision of 5%. **D)** Polysaccharide-to-lipid molar ratio determined by integrating the intensity of their respective spectral regions.

In addition to the differences in polysaccharide composition, the polysaccharide-to-lipid ratio also differed substantially between the two fractions (**Figure 4D**). The top fraction exhibited a markedly higher polysaccharide-to-lipid ratio (37-fold higher), indicating a lower relative contribution of lipid resonances in the top fraction compared to the bottom fraction (8-fold low).

### Molecular dynamics of the cell wall differs between layers and growth conditions

Next, to investigate the dynamic heterogeneity of the cell wall, we compared ^13^C-detected spectra recorded using the complementary INEPT polarization scheme, which is sensitive to highly mobile species (**Figure 5**). The resulting ^13^C spectral fingerprints differed markedly (**Figure 5A**), revealing the coexistence of rigid and mobile polysaccharide populations within the cell wall, an observation already made in previous NMR studies on the fungal cell wall (Chakraborty et al., 2021; Lamon et al., 2023). Comparison of the INEPT spectra of the two fractions revealed very similar spectral fingerprints throughout the polysaccharide region (52–110 ppm), suggesting that the composition of the highly mobile polysaccharides is largely conserved between the two mobile layers **(Figure 5B**). Due to spectral overlap and lower signal intensities, assignment of NMR signals in the mobile fraction using 1D spectroscopy was not possible.

**Figure 5.**
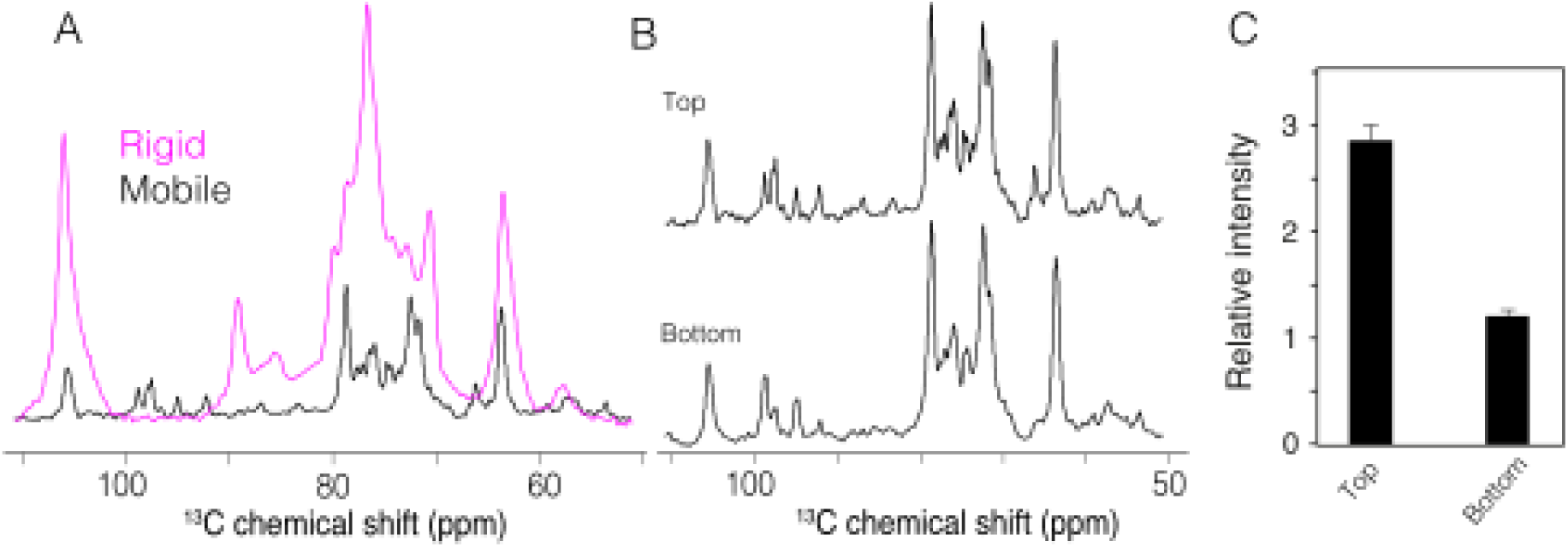
Dynamic heterogeneity of the cell wall. **A)** Spectral ^13^C fingerprints of the top fraction, recorded using two types of polarization transfers (magenta: CP and black: INEPT), revealing respectively rigid and mobile molecule of the cell wall. **B)** Comparison of the INEPT spectra of the top and bottom fractions. **C)** CP-to-INEPT intensity ratio for the top and bottom fractions.

Together, these observations indicate that the major molecular differences between the two fractions reside predominantly within the rigid molecular network. To obtain a global estimate of the relative proportion of rigid and mobile molecular components, integrated CP and INEPT signal intensities were compared (**Figure 5C**). The top fraction exhibited a CP/INEPT ratio of approximately 2.9, whereas the bottom fraction displayed a ratio of approximately 1.2, indicating a larger contribution from rigid molecular components in the top layer. Data obtained through the CP and INEPT signals analysis were combined to establish a model of the cell wall remodeling during differentiation of generative cells into skeletal cells in *T. versicolor* (**Figure 6**).

**Figure 6.**
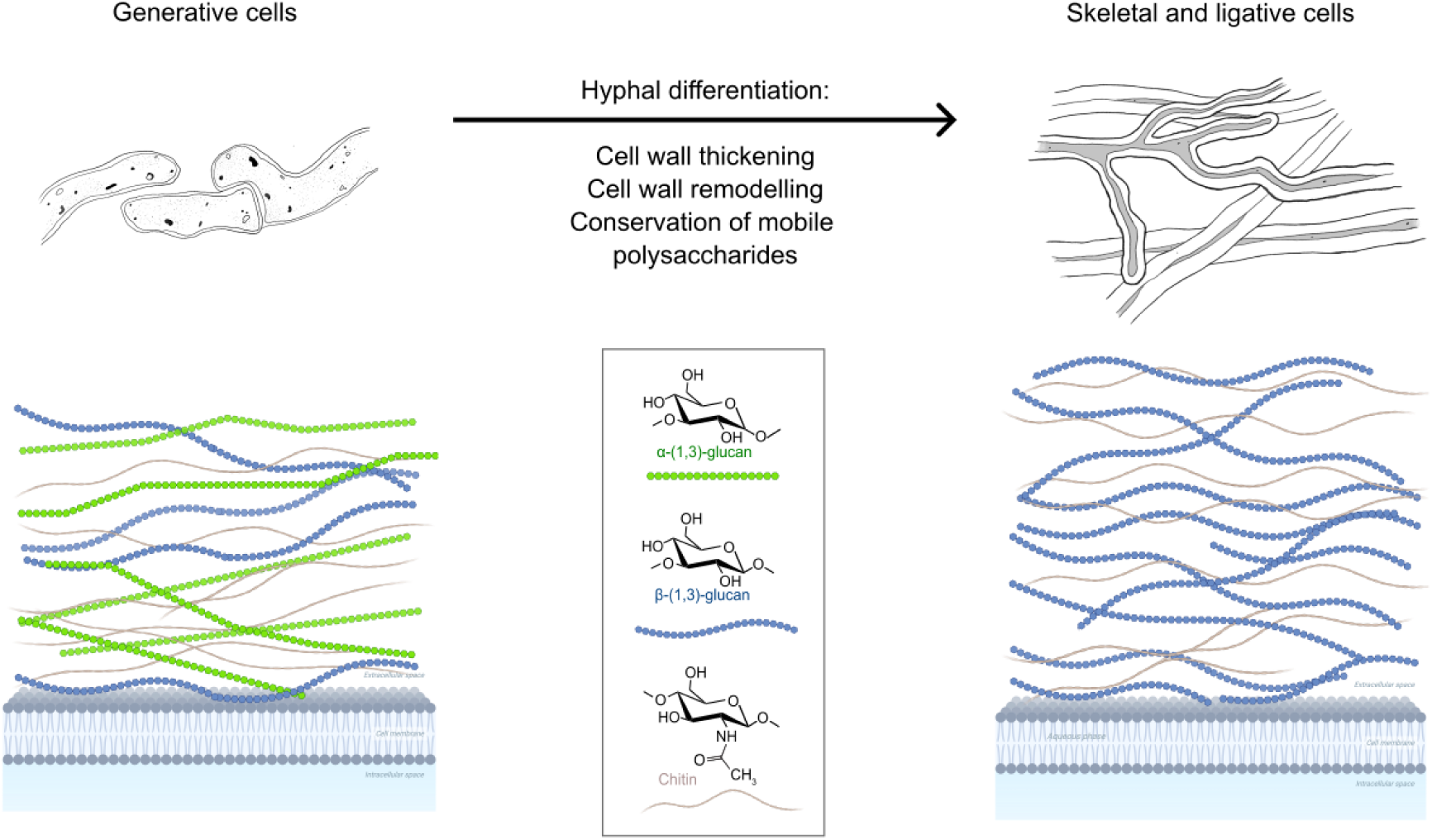
Model of the differentiation of generative cells into skeletal and ligative cells in *T. versicolor* LSSF-derived mats. Generative cells rigid components were composed of 26% β-1,3-glucan, 37% chitin and 37% α-1,3-glucan. Skeletal cells rigid components were composed of 74% β-1,3-glucan and 26% chitin with a lipids-to-polysaccharides. Mobile phases of both cell types were composed of similar polysaccharides that differ from the ones in the rigid phase, with a rigid-to-mobile ratio of 1.2 for the generative cells and of 2.9 for the skeletal cells. This study did not take into account the spatial location of the polysaccharides and their placement here is arbitrary.

Together, these observations indicate that the major molecular differences between the two fractions reside predominantly within the rigid molecular network. To obtain a global estimate of the relative proportion of rigid and mobile molecular components, integrated CP and INEPT signal intensities were compared (**Figure 5C**). The top fraction exhibited a CP/INEPT ratio of approximately 2.9, whereas the bottom fraction displayed a ratio of approximately 1.2, indicating a larger contribution from rigid molecular components in the top layer. Data obtained through the CP and INEPT signals analysis were combined to establish a model of the cell wall remodeling during differentiation of generative cells into skeletal cells in *T. versicolor* (**Figure 6**).

In addition to LSSF-derived mycelium, different tissues of a *T. versicolor* fruiting body were analyzed (**Supplementary Figures S5** and **S6**, **Supplementary Table S6**). Samples from fruiting bodies featured different cell wall composition profiles as compared to the mycelium mats, with regards to the chitin to β-(1,3)-glucan ratio. Though the ratios were different, the fact only chitin and beta-(1,3)-glucan were detected indicates these samples are more similar to the top layer of the LSSF mats than to the generative layer (**Supplementary Figures S5** and **S6**). Generative cells from a shaken liquid culture were also analyzed using NMR. Analysis of the CP signal showed that the composition of rigid components was relatively similar to that of the generative cell layer from LSSF, including the detection of α-(1,3)-glucan. However, the nature of the mobile component of the cell wall of these pellets differed from that of cells grown in LSSF, in both the bottom and top layer.

## Discussion

### Cultivation conditions determine cell type production

Microscopy of the mycelium mats allowed us to identify two layers composed of different cell types separated by a pigmented line. Similar pigmentation patterns can be observed in fruiting bodies of *Phylloporia spp.* between two different tissues (Yombiyeni et al., 2015), hinting at cell differentiation in the two layers. When cultivated in shaken liquid culture, the organism developed generative hyphae only, suggesting that the generative morphology at the bottom of the mat could be due to contact with water during the proliferation phase of LSSF. Once the surface is colonized and serves as a physical anchor, top cells grow and differentiate into skeletal and ligative hyphae.

These observations raise the question of to what extent the fermentation setup determines mycelium material quality. Mycelium leather-like materials can be produced in different fermentation setups including submerged bioreactor setups, LSSF and solid-state fermentation (Vandelook et al., 2021). Though submerged fermentation is an easily scalable process using bioreactors for biomass production, it does not seem to promote the differentiation of cells into thick-walled, semi-solid fiber-like structures, which could explain why materials produced in these setups typically result in plastic- or film-like materials rather than felt- or leather-like materials (Ahokas et al., 2026; Haneef et al., 2017).

To our knowledge, LSSF is a completely artificial growth setup that cannot be found in nature. These culture conditions might explain the significantly smaller diameter of the lab-grown skeletal cells compared with the ones found in the fruiting body. Given the significant differences in skeletal hyphae morphology under physiological *versus* artificial growth conditions, extrapolating the results from (Porter and Naleway, 2022) on the quantification of the contribution of generative, skeletal and ligative hyphae to the compressive strength of the mushroom would be an overinterpretation. The differences in skeletal hyphal diameter and cell wall composition between the mat and fruiting body do not support to the hypothesis that the organism reproduces a native tissue architecture under laboratory conditions.

### Cell type organization inffuences mechanical and physical properties

In this study, the untreated mycelium mats from *T. versicolor* cultivated in LSSF reached a UTS of 0.40 ± 0.08 MPa, an EAB of 3.90 ± 0.13% and a Young’s modulus of 27.5 ± 4.3 MPa. These values define a weak, low strain and moderately stiff material that compares to low-density rigid foams such as expanded polystyrene packaging foam and to untreated mycelium materials from other strains (Chen et al., 2024, 2015). The low EAB (∼4%) combined with the relatively high modulus indicates brittle behavior. However, the mats were deliberately not post-processed to quantify the impact of the strain’s cell morphology and organization on the material. Applying densification or plasticization steps such as heat pressing, tanning, glycerol treatment or crosslinking allows the material properties to become plastic film- or leather-like (d’Errico et al., 2025; Rovers and Wösten, 2026).

When examining individual stress-strain plots of each technical replicate, a few smaller failures can be observed before complete failure of the material. Those drops correspond to sudden failures of the generative bottom layer, following a brittle fracture behavior. After failure of the generative hyphae, the skeletal hyphae fail at higher strain following a ductile failure behavior. These observations are consistent with the SEM observations of the material: the generative bottom layer, composed of highly densely aggregated, thin hollow cell wall tubes, fails as a single film layer. On the other hand, the skeletal layer, composed of loosely aggregated, fiber-like cells with almost no lumen, behaves as a non-woven or felt-like material (Woo, 2021).

Cellular organization was consistent with the WCA, hysteresis and DVS results. This showed that the bottom side of the material, which features densely aggregated cells forming a smoother and homogeneous surface, allows water droplets to slide more easily than the top side. The top side, which is rougher and more heterogeneous, features higher adhesion consistent with a “rose petal” behavior. However, when exposed to humidity for a longer period, the density with which the cells are organized seems to play a more important role. While the skeletal layer was more hydrophobic at the droplet level, it absorbed more water at RH levels below 80%, which could be explained by its looser matrix structure allowing for a larger internal surface area for water vapor. In contrast, the generative layer, formed by densely aggregated cells, presents less surface area for vapor absorption.

### Cell wall remodeling accompanies hyphal differentiation

One of the most striking findings of this study is the extensive remodeling of the rigid cell wall accompanying the differentiation of generative hyphae into skeletal and ligative hyphae. Solid-state NMR revealed that the upper layer of the mycelium mat completely lacks detectable α-1,3-glucan, whereas this polysaccharide represents approximately one-third of the rigid cell wall in the generative bottom layer. Conversely, the skeletal/ligative fraction is composed almost exclusively of β-1,3-glucan and chitin. These results demonstrate that hyphal differentiation in *T. versicolor* is accompanied by profound changes in the molecular architecture of the cell wall rather than by simple thickening of the cell wall. Although this difference initially appears surprising, similar trends have been reported in other basidiomycetes used for biomaterial production. In *Fomes fomentarius*, another member of the Polyporaceae family, the context tissue, which consists predominantly of skeletal and ligative hyphae, also lacks detectable α-1,3-glucan and is largely composed of β-1,3-glucan (Pylkkänen et al., 2023). In contrast, tissues enriched in generative hyphae such as the crust retain α-1,3-glucan. Together, these observations suggest that the extensive remodeling of polysaccharide composition accompanying hyphal differentiation may represent a common feature among trimitic Polyporales rather than a peculiarity of *T. versicolor*.

The functional consequences of this remodeling remain to be fully established. β-1,3-glucans assemble into triple-helical structures stabilized by hydrogen bonding, providing both mechanical strength and structural flexibility to fungal cell walls (Ruiz-Herrera C Ortiz-Castellanos, 2019; Synytsya C Novák, 2013), whereas α-1,3-glucans have been associated with more compact, aggregative and supramolecular organizations (Kobayashi et al., 2017; Yoshimi et al., 2017). Our mechanical measurements are consistent with these structural differences: the α-1,3-glucan-rich generative layer fails by brittle fracture, whereas the β-1,3-glucan-rich skeletal layer displays a more ductile behavior (Togo et al., 2021). While the distinct cellular organization of the two layers likely contributes to their mechanical response, the marked differences in polysaccharide composition may represent an additional determinant of their mechanical properties. These findings also seem to align with the vapor sorption behavior with the generative layer, richer in lipids and α-1,3-glucan, took up less water at lower relative humidities than its skeletal counterpart.

Interestingly, comparison of CP and INEPT NMR spectra further revealed that the molecular differences between the two layers are primarily associated with the rigid molecular components of the cell wall. In contrast, the INEPT spectra of the two fractions were largely similar, indicating that the composition of the highly mobile polysaccharide populations is relatively conserved. This observation suggests that hyphal differentiation primarily remodels the rigid structural scaffold of the cell wall while maintaining a comparable mobile polysaccharide fraction.

Finally, comparison with fruiting body tissues and with mycelium grown under shaken liquid culture conditions demonstrated that the molecular architecture observed in LSSF is not simply equivalent to that of naturally occurring fungal tissues. Both the composition of the rigid polysaccharides and the dynamic organization of the cell wall differed substantially from those observed in the fruiting body **(Supplementary Figures S5** and **S6)**, indicating that LSSF promotes the formation of a distinct developmental state adapted to engineered mycelium materials.

### Importance of strain choice for material properties

*P. ostreatus* and *S. commune* are both saprotrophic agaricomycetes used for making biomaterials, but belong to a different order (Agaricales) than *T. versicolor* and *F. fomentarius*. In *P. ostreatus* (a monomitic fungus), even though the cell wall features all three polysaccharides (α-1,3-glucan, β-1,3-glucans and chitin), the majority of the cell wall is composed of other polysaccharides (Schiphof et al., 2026). It is possible to visualize the localization of the β-1,3-glucans in the outer cell wall (Nakazawa et al., 2024). Similarly, the cell wall of *S. commune* (a dimitic fungus) grown on agar is also composed of α-(1-3)-glucans, β-(1,3)-glucans and chitin but also numerous other sugars such as mannose and fucose (Ehren et al., 2020). These differences in composition between fungi from different orders (Agaricales vs Polyporales) within the same class (Agaricomycetes) show that beyond cell type, the composition of the cell wall is highly dependent on the fungus. It can be hypothesized that the more phylogenetic distant strains are, the more they diverge in the relative proportion and organization of their cell wall components.

Beyond phylogeny, these differences in cell wall composition and hyphal architecture can also be linked to the ecological strategies of the fruiting bodies. Many polypores (such as *F. fomentarius* and *Ganoderma lucidum*) form perennial, long-lived fruiting bodies that must endure years of environmental stress, desiccation, and mechanical wear. This long-term structural demand may have driven the evolutionary selection for heavily thickened, lignified cell walls and robust dimitic or trimitic hyphal systems, sometimes with intermediate structures like skeletal-binding hyphae. Exploring these specific structural adaptations could provide valuable insights for engineering PMMs. In their review, (Rovers and Wösten, 2026) established that the most important contributor to the mechanical strength of PMMs is the fungal strain, and that basidiomycetes represent the majority of strains used for making PMMs. In their list of strains used for making PMMs, a substantial proportion are dimitic or trimitic, suggesting that thickened cell walls and cell length may play an important role in mechanical properties.

### Future perspectives

Combining the findings of this paper with literature on biotechnological applications of filamentous fungi, adapting the fermentation setup to promote the growth of specific hyphal types appears to be an essential process optimization strategy. LSSF appears as a relevant fermentation technique to promote the production of differentiated fungal cells, which can then be valorized for mycelium materials with tuned physical and mechanical properties. In addition to mycelium materials applications, skeletal cells of *T. versicolor* could be valorized for their chitin- and glucan-rich cell walls by spinning into fibers for textile applications, nanopapers for water treatments or heavy metals absorption (Janesch et al., 2020; Svensson et al., 2024). With a strong oxidative enzymatic machinery coupled to cell wall biosorption capability, *T. versicolor* is an organism of choice for bioremediation processes against environmental pollutants. This capacity is used for the remediation of synthetic dyes from textile industry, but also for paper pulp whitening, removal of pesticides, pharmaceuticals, endocrine-disrupting compounds, polycyclic aromatic hydrocarbons (PAHs), industrial waste products from waste waters (Zhuo et al., 2021), lignocellulosic biomass conversion, biofuels and biorefinery approaches (Tišma et al., 2021). While biosorption activities may be enhanced by thick-walled skeletal cells, the dual generative/skeletal composition of the LSSF grown mycelium mats may make them suitable for applications requiring functional materials combining mechanically performant and metabolically active cells.

## Conclusions

This study analyzed the cellular organization of *T.* versicolor grown in LSSF and revealed that the material is not a homogeneous sheet of biomass but a bilayer composed of generative cells at the liquid interface and of skeletal and ligative cells at the air interface. Each layer carries a distinct identity at every scale examined: the generative layer is composed of thin-walled cells rich in α-1,3-glucan, densely packed. Together, these cells form a hydrophilic surface that fails by brittle fracture as a continuous film. On the other hand, the skeletal-ligative layer is composed of semi-solid fiber-like cells depleted of α-1,3-glucan and rich in β-1,3-glucan and chitin. This layer is also more lipid-rich and features a larger rigid fraction than the generative cells. Together, these skeletal-ligative cells assemble like a non-woven material and create a hydrophobic and strongly water-pinning surface that fails in a ductile manner. Hyphal differentiation in this fungus therefore involves remodeling of the molecular cell wall architecture rather than simple cell wall thickening, and the mechanical and surface behavior of the material can be associated with that remodeling and cellular organization.

Equally important is what this material is not. Skeletal hyphae formed in liquid state surface fermentation are about half the diameter of those in the fruiting body, and both the rigid polysaccharide composition and the cell wall dynamics of the cultivated tissues differ from those of fruiting-body context, crust and pore tissues. LSSF thus appears to generate a distinct developmental state of the fungus.

## List of abbreviations

LSSF: Liquid State Surface Fermentation
PMM: Pure Mycelium Materials
CYM: Complete Yeast Medium
EAB: Elongation At Break (%)
UTS: Ultimate Tensile Stress (MPa)
CP: Cross Polarization
INEPT: Insensitive Nuclei Enhanced by Polarization Transfer
E: Young’s modulus (MPa)
WCA: Water Contact Angle (°)
DVS: Dynamic Vapor Sorption
RH: Relative Humidity

## Declarations

### Ethics approval and consent to participate

Not applicable.

### Consent for publication

Not applicable.

### Availability of data and materials

The datasets generated and analyzed during the current study are available in the Supplementary Materials or in the Zenodo repository, https://doi.org/10.5281/zenodo.24116122, with the exception of NMR data, which are available upon request to AL.

### Authors’ contributions

AOS, MA and EP conceptualized the research. AOS and AL wrote the original draft and made the figures. AOS performed all experiments besides NMR and DVS. NMR was performed by AL and AE. SV performed the DVS. PA produced the samples used for part of the microscopy pictures. JB and NVB contributed in conceptualizing the mechanical properties analysis and supervised the tensile strength testing. SV, LD and KDC contributed in defining the material surface characterization strategy. MA helped with the microscopic analysis. All authors reviewed and edited the manuscript.

## Acknowledgements

This work has benefited from the Biophysical and Structural Chemistry Platform at IECB, CNRS UAR3033, INSERM US001, University of Bordeaux. The authors would like to acknowledge Priya Laha for the help in making picture of figure 1E, Anouk Verstuyft for her render of the mycelium mat in the graphical abstract and Juliana Masis Solano for making Supplementary Figure S2. The graphical abstract and Figure 6 were made partially using Biorender.com.

## Authors’ information

Annah-Ọọl ládé Ṣàngósànyà earned her MEng in Biological engineering from the National Institute of Applied Sciences (INSA) of Toulouse (FR) in 2021, specializing in microbiology and synthetic biology. She then pursued the Fabricademy: Textile and Technology academy postgraduate from the Universitat Politecnica de Catalunya (ES), an intensive applied research program focused on new technologies applied to the textile industry. After receiving the Young Scientist Award (Lenzing) grant rewarding her postgraduate thesis, she worked for a year as an independent researcher on textile degradation using mycelium. Currently, as a PhD candidate at Vrije Universiteit Brussel (VUB), AOS focuses on exploring biodiversity to understand the cellular structures making certain fungal strains suitable for mycelium materials and on their potential for degrading textile waste.

## Footnotes

Not applicable.

## References

Ahokas P, Arias-Barrantes M, Valtonen A, Kärkkäinen E, Kunnari V, Vikman M, et al. Production of Mycelium-Based Nonwoven Fabrics via Submerged Fermentation. ACS Appl Bio Mater 2026. 10.1021/acsabm.6c00471.

Aiduang W, Jatuwong K, Luangharn T, Jinanukul P, Thamjaree W, Teeraphantuvat T, et al. A Review Delving into the Factors Influencing Mycelium-Based Green Composites (MBCs) Production and Their Properties for Long-Term Sustainability Targets. Biomimetics 2024;9:337. 10.3390/biomimetics9060337.

Aquino M, Rugolo M, Robledo G, Kuhar F. Evaluation of mycelium composite materials produced by five Patagonian fungal species. Maderas Ciencia y Tecnología 2022;24. 10.4067/S0718-221X2022000100435.

Bayramoglu G. Biosorption of heavy metal ions on immobilized white-rot fungus Trametes versicolor. J Hazard Mater 2003;101:285–300. 10.1016/S0304-3894(03)00178-X.

Boujelben R, Ellouze M, Tóran MJ, Blánquez P, Sayadi S. Mycoremediation of Tunisian tannery wastewater under non-sterile conditions using Trametes versicolor: live and dead biomasses. Biomass Convers Biorefin 2024;14:299–312. 10.1007/s13399-022-02328-0.

Brandon Matheny P, Wang Z, Binder M, Curtis JM, Lim YW, Henrik Nilsson R, et al. Contributions of rpb2 and tef1 to the phylogeny of mushrooms and allies (Basidiomycota, Fungi). Mol Phylogenet Evol 2007;43:430–51. 10.1016/j.ympev.2006.08.024.

Cartabia M, Girometta CE, Milanese C, Baiguera RM, Buratti S, Branciforti DS, et al. Collection and Characterization of Wood Decay Fungal Strains for Developing Pure Mycelium Mats. Journal of Fungi 2021;7:1008. 10.3390/jof7121008.

Cengiz G, Aytar P, Şam M, Çabuk A. Removal of Reactive Dyes using Magnetically Separable *Trametes versicolor* Cells as a New Composite Biosorbent. Sep Sci Technol 2014;49:1860–71. 10.1080/01496395.2014.903496.

Chakraborty A, Fernando LD, Fang W, Dickwella Widanage MC, Wei P, Jin C, et al. A molecular vision of fungal cell wall organization by functional genomics and solid-state NMR. Nat Commun 2021;12:6346. 10.1038/s41467-021-26749-z.

Chen H, Klemm S, Dönitz AG, Ou Y, Schmidt B, Fleck C, et al. Tailoring the Mechanical Properties of Fungal Mycelium Mats with Material Extrusion Additive Manufacturing of PHBH and PLA Biopolymers. ACS Omega 2024;9:49609–17. 10.1021/acsomega.4c07661.

Chen W, Hao H, Hughes D, Shi Y, Cui J, Li Z-X. Static and dynamic mechanical properties of expanded polystyrene. Mater Des 2015;69:170–80. 10.1016/j.matdes.2014.12.024.

Corner EJH. The Fruit-body of Polystictus xanthopus Fr. Ann Bot 1932a;46:71–111.

Corner EJH. A Fomes with two systems of hyphae. Transactions of the British Mycological Society 1932b;17:51–81.

d’Errico A, van den Brandhof JG, Bogomolova A, Wösten HAB. Strongest untreated mycelium materials produced by Schizophyllum commune dikaryons. World J Microbiol Biotechnol 2025;41:403. 10.1007/s11274-025-04582-6.

Ehren HL, Appels FVW, Houben K, Renault MAM, Wösten HAB, Baldus M. Characterization of the cell wall of a mushroom forming fungus at atomic resolution using solid-state NMR spectroscopy. The Cell Surface 2020;6:100046. 10.1016/j.tcsw.2020.100046.

Elgarahy AM, Elwakeel KZ, Mohammad SH, Elshoubaky GA. A critical review of biosorption of dyes, heavy metals and metalloids from wastewater as an efficient and green process. Clean Eng Technol 2021;4:100209. 10.1016/j.clet.2021.100209.

Gallagher JAL, Redmond JN, Asgari M, Bradshaw AJ, Dentinger BTM, Naleway SE. Generative hyphal stiffness and cell wall thickening in fungi. Journal of Materials Research and Technology 2025;37:3105–14. 10.1016/j.jmrt.2025.06.209.

Gandia A, van den Brandhof JG, Appels FVW, Jones MP. Flexible Fungal Materials: Shaping the Future. Trends Biotechnol 2021;39:1321–31. 10.1016/j.tibtech.2021.03.002.

Habtemariam S. Trametes versicolor (Synn. Coriolus versicolor) Polysaccharides in Cancer Therapy: Targets and Efficacy. Biomedicines 2020;8:135. 10.3390/biomedicines8050135.

Haneef M, Ceseracciu L, Canale C, Bayer IS, Heredia-Guerrero JA, Athanassiou A. Advanced Materials From Fungal Mycelium: Fabrication and Tuning of Physical Properties. Sci Rep 2017;7:41292. 10.1038/srep41292.

Janesch J, Jones M, Bacher M, Kontturi E, Bismarck A, Mautner A. Mushroom-derived chitosan-glucan nanopaper filters for the treatment of water. React Funct Polym 2020;146:104428. 10.1016/j.reactfunctpolym.2019.104428.

Jones M, Gandia A, John S, Bismarck A. Leather-like material biofabrication using fungi. Nat Sustain 2021;4:9–16. 10.1038/s41893-020-00606-1.

Kleijburg FEL, Schunselaar EM, Safeer AA, Bishoyi AK, Baldus M, Wösten HAB. Functions of the cell wall polysaccharide schizophyllan during vegetative growth of Schizophyllum commune. The Cell Surface 2026;15:100167. 10.1016/j.tcsw.2025.100167.

Kobayashi K, Hasegawa T, Kusumi R, Kimura S, Yoshida M, Sugiyama J, et al. Characterization of crystalline linear (1 → 3)-α-d-glucan synthesized in vitro. Carbohydr Polym 2017;177:341–6. 10.1016/j.carbpol.2017.09.003.

Lamon G, Lends A, Valsecchi I, Wong SSW, Duprès V, Lafont F, et al. Solid-state NMR molecular snapshots of *Aspergillus fumigatus* cell wall architecture during a conidial morphotype transition. Proceedings of the National Academy of Sciences 2023;120. 10.1073/pnas.2212003120.

Lodi RS, Dong X, Wang X, Han Y, Liang X, Peng C, et al. Current research on the medical importance of Trametes species. Fungal Biol Rev 2025;51:100413. 10.1016/j.fbr.2025.100413.

Nakazawa T, Kawauchi M, Otsuka Y, Han J, Koshi D, Schiphof K, et al. Pleurotus ostreatus as a model mushroom in genetics, cell biology, and material sciences. Appl Microbiol Biotechnol 2024;108:217. 10.1007/s00253-024-13034-4.

Porter DL, Naleway SE. Hyphal systems and their effect on the mechanical properties of fungal sporocarps. Acta Biomater 2022;145:272–82. 10.1016/J.ACTBIO.2022.04.011.

Pylkkänen R, Werner D, Bishoyi A, Weil D, Scoppola E, Wagermaier W, et al. The complex structure of Fomes fomentarius represents an architectural design for high-performance ultralightweight materials. Sci Adv 2023;9. 10.1126/SCIADV.ADE5417/SUPPL_FILE/SCIADV.ADE5417_MOVIES_S1_TO_S9.ZIP.

Rea I, Giardina P, Longobardi S, De Stefano L. Protein-modified porous silicon films for biomedical applications. Porous Silicon for Biomedical Applications, Elsevier; 2014, p. 104–28. 10.1533/9780857097156.1.104.

Rehner SA, Buckley E. A Beauveria phylogeny inferred from nuclear ITS and EF1-sequences: evidence for cryptic diversification and links to Cordyceps teleomorphs. Mycologia 2005;97:84–98. 10.3852/mycologia.97.1.84.

Romero-Cedillo RK, Robledo-Leal ER, Aguilar-Marcelino L, Acosta-Urdapilleta Ma de L, Téllez-Téllez M. Production of Mycelium Mats for Textile Applications. Journal of Fungi 2025;11:700. 10.3390/jof11100700.

Rovers ICA, Wösten HAB. Fungal diversity used to produce mycelium materials. Mater Today Commun 2026;52:115164. 10.1016/j.mtcomm.2026.115164.

Ruiz-Herrera J, Ortiz-Castellanos L. Cell wall glucans of fungi. A review. The Cell Surface 2019;5:100022. 10.1016/j.tcsw.2019.100022.

Ryvarden Leif, Decock Cony, Mossebo D., Masuka AJ., Melo I. Poroid fungi of Africa. Fungiflora; 2022.

Ryvarden Leif, Melo I. Poroid fungi of Europe. 1st ed. Fungiflora; 2014.

Schiphof K, Hitaka A, Kawauchi M, Nakazawa T, Honda Y. Functional analysis of the class III chitin synthase genes in the filamentous basidiomycete Pleurotus ostreatus. Fungal Genetics and Biology 2026;184:104081. 10.1016/j.fgb.2026.104081.

Silm M, Jiang Q, Kisand A, Bismarck A, Jones MP. Transforming textile waste into materials using fungi. Resour Conserv Recycl 2026;228:108812. 10.1016/j.resconrec.2026.108812.

Svensson SE, Wijayarathna ERKB, Kalita NK, Hakkarainen M, Zamani A. Development of hydrogels from cell wall of Aspergillus oryzae containing chitin-glucan and wet spinning to monofilaments. Int J Biol Macromol 2024;278:134285. 10.1016/j.ijbiomac.2024.134285.

Synytsya A, Novák M. Structural diversity of fungal glucans. Carbohydr Polym 2013;92:792–809. 10.1016/j.carbpol.2012.09.077.

Tišma M, Žnidaršič-Plazl P, Šelo G, Tolj I, Šperanda M, Bucić-Kojić A, et al. Trametes versicolor in lignocellulose-based bioeconomy: State of the art, challenges and opportunities. Bioresour Technol 2021;330:124997. 10.1016/j.biortech.2021.124997.

Togo A, Suzuki S, Kimura S, Iwata T. Wet Spinning and Structure Analysis of α-1,3-Glucan Regenerated Fibers. ACS Appl Polym Mater 2021;3:2063–9. 10.1021/acsapm.1c00114.

Vandelook S, Bassleer B, Elsacker E, Peeters E. Effects of Orange Peel Extract on Laccase Activity and Gene Expression in Trametes versicolor. Journal of Fungi 2024;10:370. 10.3390/jof10060370.

Vandelook S, Elsacker E, Van Wylick A, De Laet L, Peeters E. Current state and future prospects of pure mycelium materials. Fungal Biol Biotechnol 2021;8. 10.1186/s40694-021-00128-1.

White TJ, Bruns T, Lee S, Taylor J. AMPLIFICATION AND DIRECT SEQUENCING OF FUNGAL RIBOSOMAL RNA GENES FOR PHYLOGENETICS. PCR Protocols, Elsevier; 1990, p. 315–22. 10.1016/B978-0-12-372180-8.50042-1.

Woo S. Mechanical System Failure. Design of Mechanical Systems Based on Statistics, First Edition. | Boca Raton : CRC Press, 2021. |: CRC Press; 2021, p. 197–233. 10.1201/9780429022050-7.

Yalçınkaya Y, Soysal L, Denizli A, Arıca MY, Bektaş S, Genç Ö. Biosorption of cadmium from aquatic systems by carboxymethylcellulose and immobilized Trametes versicolor. Hydrometallurgy 2002;63:31–40. 10.1016/S0304-386X(01)00200-6.

Yombiyeni P, Balezi A, Amalfi M, Decock C. Hymenochaetaceae from the Guineo-Congolian rainforest: three new species of *Phylloporia* based on morphological, DNA sequences and ecological data. Mycologia 2015;107:996–1011. 10.3852/14-298.

Yoshimi A, Miyazawa K, Abe K. Function and Biosynthesis of Cell Wall α-1,3-Glucan in Fungi. Journal of Fungi 2017;3:63. 10.3390/jof3040063.

Zhuo R, Fan F, Verlicchi P. A comprehensive insight into the application of white rot fungi and their lignocellulolytic enzymes in the removal of organic pollutants. Science of the Total Environment 2021;778:146132. 10.1016/j.scitotenv.2021.146132.

