## Supplementary Figures and Tables for "Hyphal differentiation and cell wall remodeling in *Trametes versicolor* underlie a layered architecture in pure mycelium materials"

### Supplementary Materials

**Cell differentiation and organization in leather-like mycelium materials: a deep dive into *Trametes versicolor* hyphal system**

Authors: Annah-Ololade Sangosanya*[1], Antoine Loquet[2], Agathe Ecoutin[2], Sofie Verschraegen[3], Phebe Arnauts[1], Joost Brancart[4] , Niko Van den Brande[4], Lode Daelemans[3], Karen De Clerck[3], Mario Amalfi[5,6], Eveline Peeters *[1] .

[1] Research group of Microbiology, Department of Bioengineering Sciences, Vrije Universiteit Brussel, Pleinlaan 2, B-1050 Brussels, Belgium.
[2] Université Bordeaux, CNRS, Bordeaux INP, CBMN, UMR 5248, IECB, F-33600 Pessac, France.
[3] Centre for Textile Science and Engineering (CTSE), Department of Materials, Textiles and Chemical Engineering (MaTCh), Ghent University, B-9052 Zwijnaarde (Ghent), Belgium.
[4] Sustainable Materials Engineering, Vrije Universiteit Brussel, Pleinlaan 2, B-1050 Brussels, Belgium.
[5] Meise Botanic Garden, Research Department, Nieuwlaan 32, B-1860 Meise, Belgium.
[6] Fédération Wallonie-Bruxelles, Service Général de l'Enseignement Supérieur et de la Recherche Scientifique, Brussels, Belgium.

#### Supplementary figures

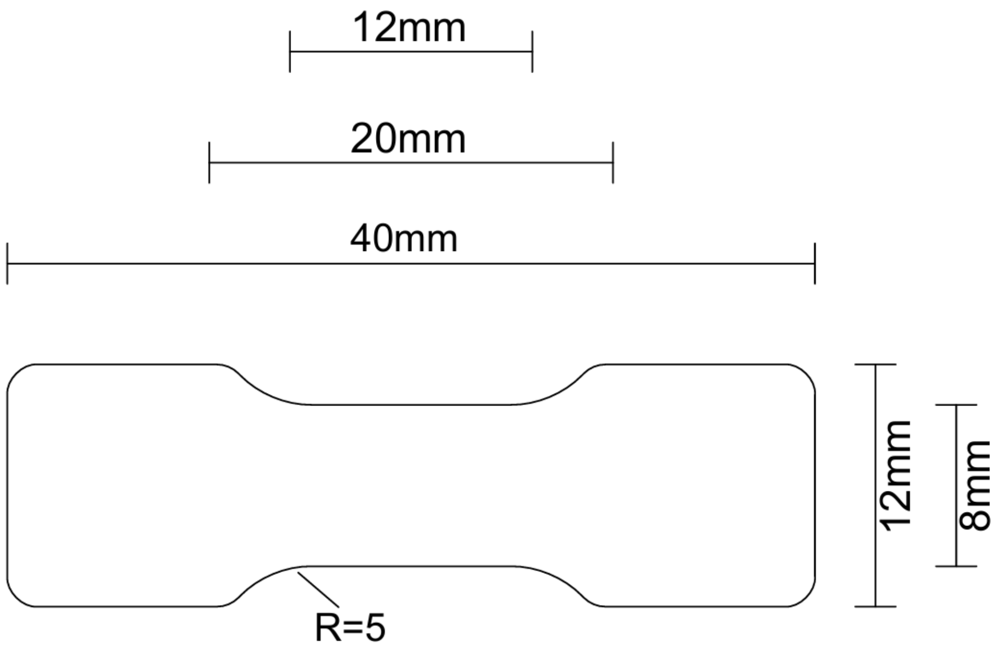


***Supplementary Figure S1:** Specifications of the custom-made dog bone shape cutting die used for tensile strength testing. All radiuses are 1.5 unless specified otherwise.*

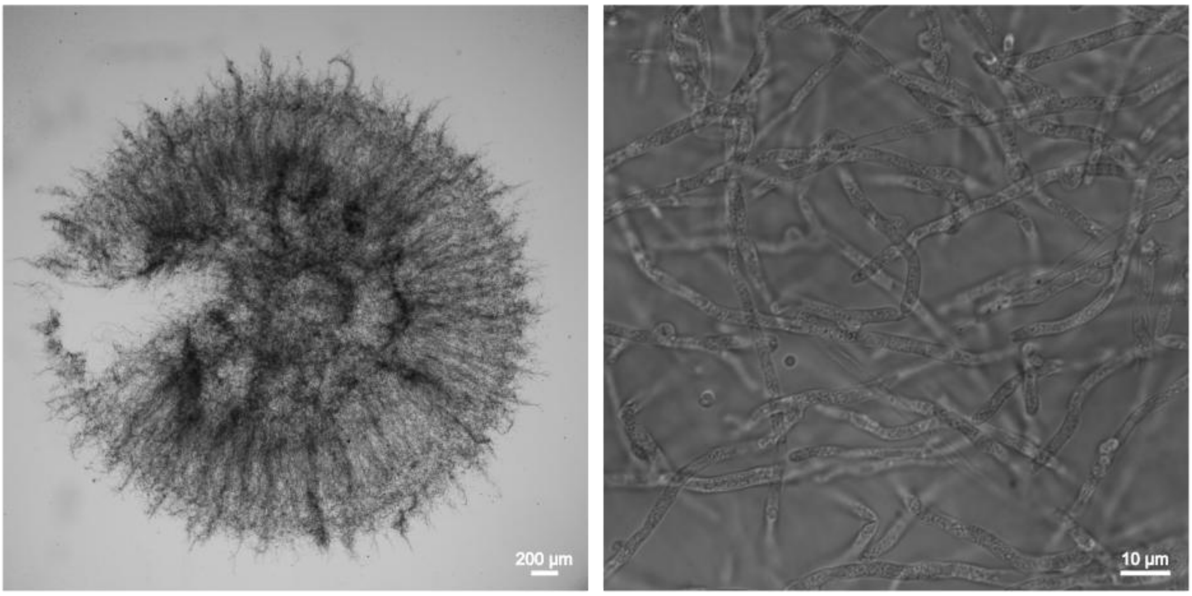


***Supplementary Figure S2****: generative hyphae (right) produced in mycelium pellets (left).*

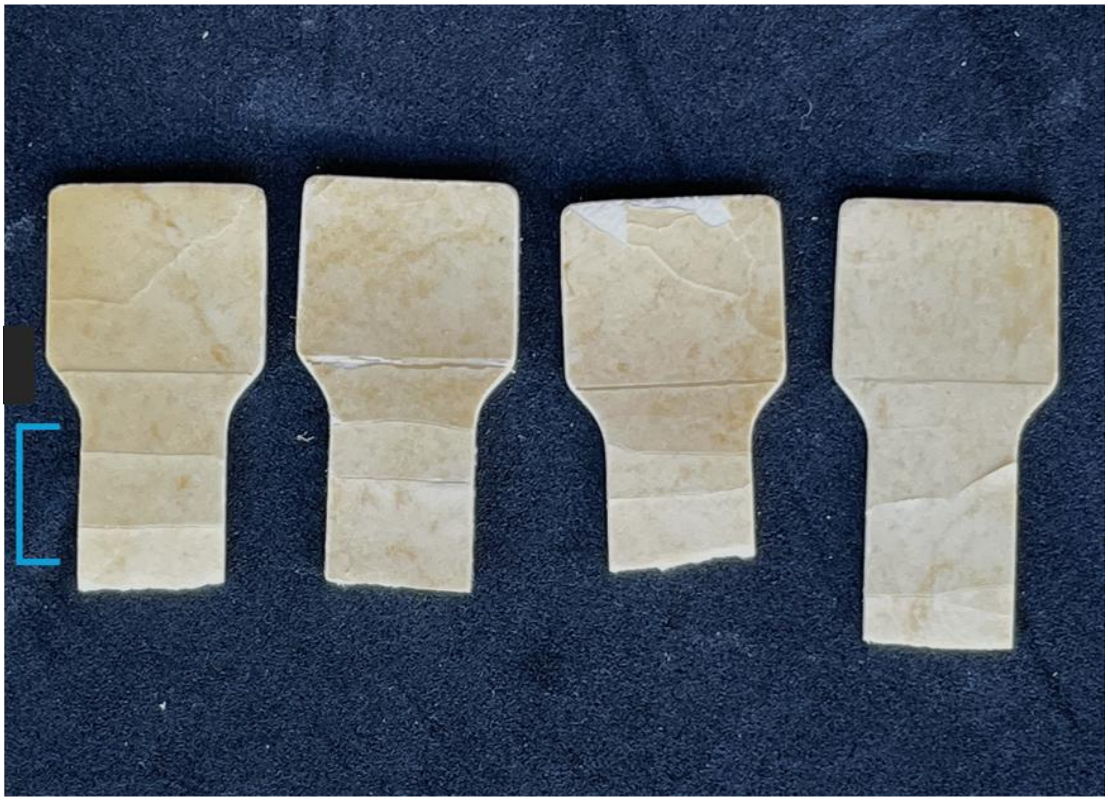

***Supplementary Figure S3: tensile strength tested samples showing generative layer failure (blue bracket).***
*Image by Juliana Masis Solano.*

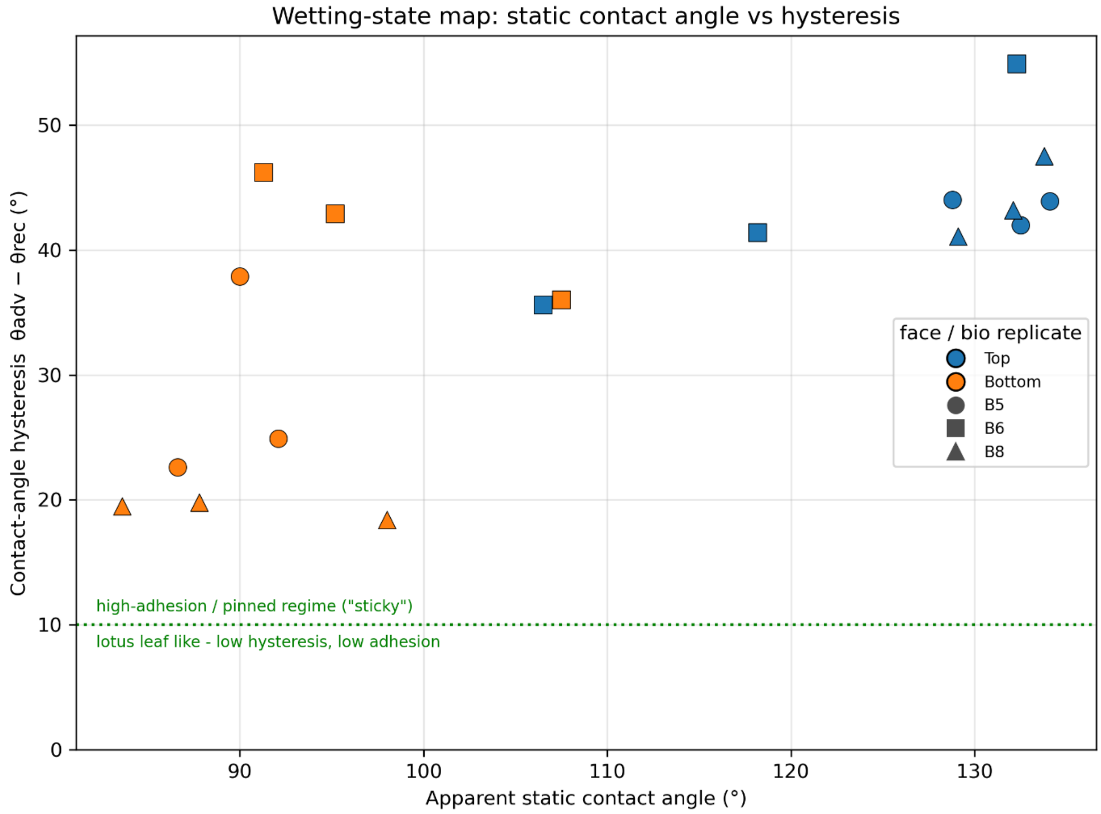


***Supplementary Figure S4: Wetting state map of the top vs bottom layer of the mat.*** *In this experiment, the surface was tilted before the droplet could be fully absorbed by the generative layer on the bottom side. The segregated clustering of the bottom layer measurements indicates that this layer slides the water droplets more easily than the top side.*

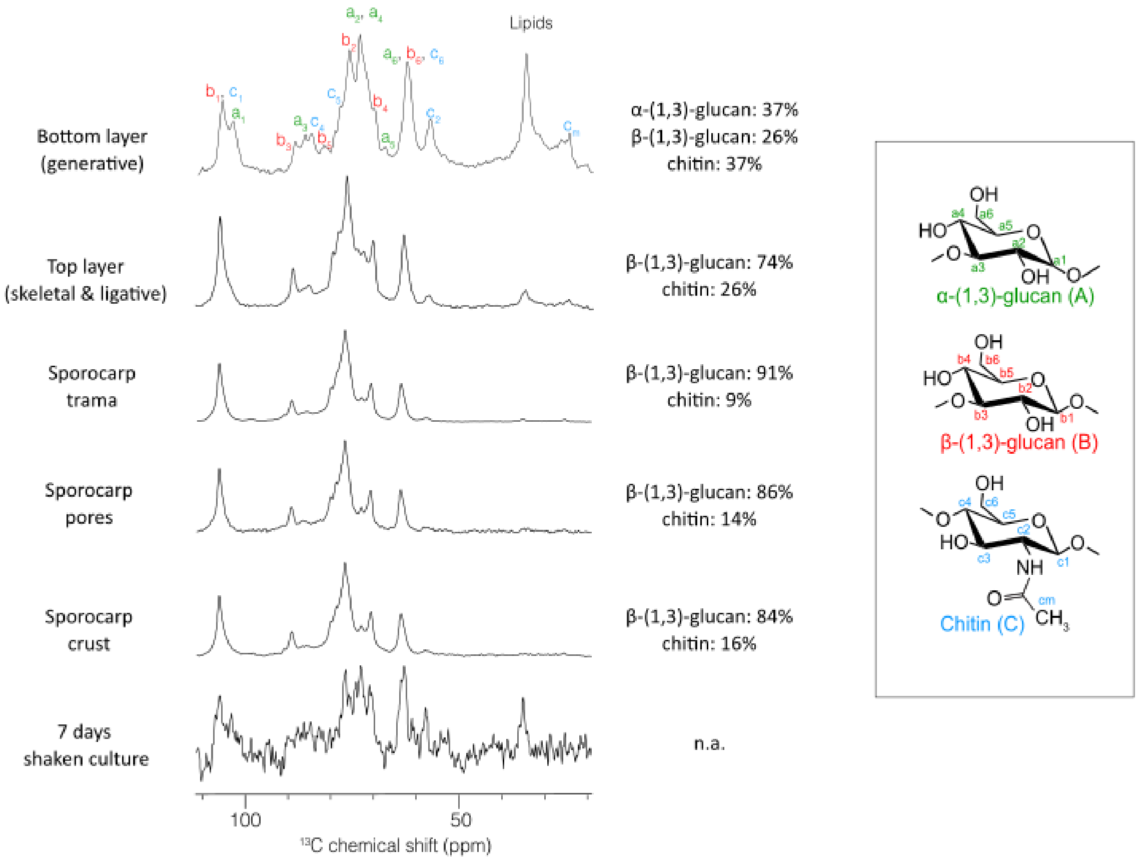


***Supplementary Figure S5: Structural analysis of the rigid components (CP signal) of the cell wall of LSSF samples, fruiting body samples and LSSF sample.*** *For the 7days submerged fermentation, the CP signal was too low to produce a reliable determination of the ratios of alpha-1,3-glucan, beta-1,3-glucan and chitin. Nevertheless, a visual comparison indicates that it looks similar to the CP of the bottom fraction. In this sample, alpha-1,3-glucan was detected. Stucture of the different polysaccharides and the resonance of each carbon are indicated with the numbered and coloured carbons. Comparing the CP signal to the INEPT signal and calculating their ratio was used to reveal the cell wall dynamic in each tissues of the fruiting body (Additional table 6).*

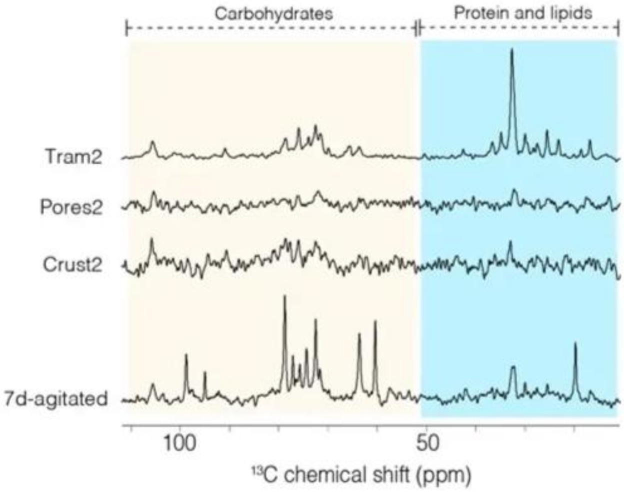


***Supplementary Figure S6: Fingerprints of the mobile component of the cell wall (INEPT signals).*** *Diving into the INEPT signals, it is observable that the INEPT spectral fingerprint of the Trama looks quite similar to what has been observed for the INEPT data on the top / bottom fractions. "7d-agitated" has a different fingerprint, suggesting the nature of the polysaccharides from the mobile component are not the same as compared to the bottom/top fractions.*

#### Supplementary tables

***Supplementary Table S1: Hyphal diameters of generative and skeletal hyphae in Trametes versicolor fruiting body and LSSF mycelium mat tissues.***

| Tissue | Hyphal type | *n* measurements | *n* images | Mean ± SD (µm) | Median (µm) |
|---|---|---|---|---|---|
| **Fruiting body** | Generative | 37 | 3 | 2.39 ± 0.51 | 2.43 |
| **Fruiting body** | Skeletal | 93 | 5 | 5.10 ± 1.05 | 4.90 |
| **Mycelium mat (LSSF)** | Generative | 70 | 5 | 2.43 ± 0.42 | 2.40 |
| **Mycelium mat (LSSF)** | Skeletal | 65 | 3 | 2.33 ± 0.50 | 2.17 |

***Supplementary Table S2: Pairwise comparisons of estimated marginal hyphal diameters*** *(linear mixed model, Holm-adjusted). Hyphal diameters (N = 265 measurements from 16 images) were analysed with a linear mixed model with tissue/hyphal-type group as fixed effect and image as random effect (Diameter ~ Group + (1|Image), maximum likelihood). The group effect was significant (likelihood-ratio test: $\chi^2 = 49.13$, df = 3, $p = 1.2 \times 10^{-10}$). Between-image variance was 0.041 µm$^2$, residual variance 0.483 µm$^2$. Pairwise contrasts were Holm-corrected for multiple testing. FB = fruiting body; Mat = LSSF mycelium mat.*

| Comparison | Δ mean (µm) | Holm-adjusted *p* | Significant |
|---|---|---|---|
| **FB Skeletal – FB Generative** | +2.70 | < 0.001 | Yes |
| **FB Skeletal – Mat Generative** | +2.73 | < 0.001 | Yes |
| **FB Skeletal – Mat Skeletal** | +2.69 | < 0.001 | Yes |
| **Mat Generative – FB Generative** | +0.03 | 1.00 | No |
| **Mat Skeletal – FB Generative** | −0.01 | 1.00 | No |
| **Mat Skeletal – Mat Generative** | −0.04 | 0.90 | No |

***Supplementary Table S3: UTS, EAB and Young's modulus of the measured samples.***

| Biological replicate | UTS mean (MPa) | UTS SD (MPa) | EAB mean (%) | EAB SD (%) | E mean (MPa) | E SD (MPa) |
|---|---|---|---|---|---|---|
| **Bio_1** | 0.348013 | 0.089663 | 4.023.517 | 0.732711 | 25.951.655 | 4.961.862 |
| **Bio_2** | 0.369332 | 0.150221 | 3.894.070 | 1.856.584 | 24.246.282 | 17.572.868 |
| **Bio_3** | 0.497262 | 0.155871 | 3.773.267 | 1.164.321 | 32.388.169 | 6.933.753 |
| **Grand mean** | 0.404869 | 0.080722 | 3.896.951 | 0.125150 | 27.528.702 | 4.293.936 |

***Supplementary Table S4: Apparent water contact angle of the top (skeletal-ligative) and bottom (generative) faces of Trametes versicolor LSSF mycelium mats, 30 s and 5 min after droplet deposition****. Each biological replicate is the mean of 5 technical replicates; mean ± SD is calculated across the 3 biological replicates. Values are 1-second time-binned apparent contact angles. Droplets that were fully absorbed by the material before the end of the acquisition were assigned a contact angle of 0°, as complete absorption represents total wetting; absorption occurred between 37 s and 294 s and only on the bottom face. Surfaces below 90° are conventionally described as hydrophilic and above 90° as hydrophobic.*

| Face | Time after deposition | *n* biological replicates | Mean ± SD (°) | Wetting regime |
|---|---|---|---|---|
| **Top** | 30 s | 3 | 135.7 ± 5.1 | Hydrophobic |
| **Top** | 5 min | 3 | 134.3 ± 5.4 | Hydrophobic |
| **Bottom** | 30 s | 3 | 90.2 ± 19.1 | at the hydrophilic/hydrophobic boundary |
| **Bottom** | 5 min | 3 | 56.5 ± 51.0 | Hydrophylic (6 technical replicates fully absorbed the droplet before the end of the experiment). |

***Supplementary Table S5: Pairwise comparisons of apparent water contact angle****. Two-tailed paired Student's t-test on biological replicate means, with technical replicates averaged within each biological replicate. Significant values ($p < 0.05$) in bold.*

| Comparison | *n* pairs | *t* | df | *p* |
|---|---|---|---|---|
| **Top, 30 s vs 5 min** | 3 | 1.347 | 2 | 0.310 |
| **Bottom, 30 s vs 5 min** | 3 | 1.773 | 2 | 0.218 |
| **Top vs Bottom, 30 s** | 3 | 5.189 | 2 | **0.035** |
| **Top vs Bottom, 5 min** | 3 | 2.878 | 2 | 0.102 |

***Supplementary Table S6:* *Cell wall dynamics.*** *Comparison of CP-to-INEPT ratio, calculated in the 50-110 region. For the pores, the INEPT signal was extremely low, indicating a highly rigid sample. Cells produced in submerged fermentation (generative cells as observed in Supp figure 2B) had a ratio comparable to the bottom layer of generative cells, which corresponds to a lower rigidity than the of skeletal and ligative cells. The crust samples and Trama samples featured a very high ratio (i.e. very weak INEPT signal), suggesting highly rigid samples. The pores had almost no INEPT signal, so it's very rigid (ratio can't be reliably computed but it should be very high).*

| | Bottom layer (generative) | Top layer (skeletal and ligative) | Sporocarp: trama | Sporocarp: crust | Sporocarp: pores | 7 days shaken culture |
|---|---|---|---|---|---|---|
| **CP/INEPT signal intensity** | 1,21 | 2,86 | 1,29 | 24,5 | n. a. (very high) | ~ 0,71 |
